# A haplotype-resolved, near telomere-to-telomere genome assembly of the desiccation-tolerant plant *Vellozia tubiflora*

**DOI:** 10.64898/2026.10.22.761125

**Authors:** Francine A. N. de Almeida, Nicholas A. Allsing, Marina P. Marone, Mariana F. Cavalheiro, Lucas E. C. Canesin, Merly Escalona, Oanh H. Nguyen, Ruta Sahasrabudhe, Diego M. Riaño-Pachón, Stephen Richards, Harris A. Lewin, Paulo Arruda, Todd P. Michael, Ricardo A. Dante, Isabel R. Gerhardt

**Affiliations:** Centro de Pesquisa em Genômica Aplicada às Mudanças Climáticas (GCCRC), Universidade Estadual de Campinas, Campinas, SP, 13083-875, Brazil; Centro de Biologia Molecular e Engenharia Genética, Universidade Estadual de Campinas, Campinas, SP, 13083-875, Brazil; Plant Molecular and Cellular Biology, Salk Institute for Biological Sciences, La Jolla, CA, 92037, USA; Embrapa Agricultura Digital, Campinas, SP, 13083-886, Brazil; Departamento de Genética, Evolução, Microbiologia e Imunologia, Instituto de Biologia, Universidade Estadual de Campinas, Campinas, SP, 13083-970, Brazil; Institute for Systems Genomics, University of Connecticut, Storrs, CT, 06269, USA; Department of Biomolecular Engineering, University of California, Santa Cruz, Santa Cruz, CA, 95064, USA; DNA Technologies and Expression Analysis Core Laboratory, Genome Center, University of California, Davis, CA, 95616, USA; Centro de Energia Nuclear na Agricultura, Universidade de São Paulo, Piracicaba, SP, 13400-970, Brazil; Department of Entomology and Nematology, University of California, Davis, Davis, CA, 95616, USA; Julie Ann Wrigley Global Futures Laboratory, Walton Center for Planetary Health, Arizona State University, Tempe, AZ, 85287 USA; Department of Cell and Developmental Biology, School of Biological Sciences, University of California, San Diego, La Jolla, CA, 92093, USA; Center for Marine Biotechnology and Biomedicine, Scripps Institute of Oceanography, University of California, San Diego, La Jolla, CA, 92037, USA; Department of Science and Conservation, San Diego Botanical Garden, Encinitas, CA, 92024, USA

## Abstract

Vegetative desiccation tolerance allows plants to survive the loss of up to 95% of their cellular water content and recover upon rehydration. The family Velloziaceae contains the largest number of desiccation-tolerant species among flowering plants, yet genomic resources for this lineage remain extremely limited. Here we report the first reference genome of the genus *Vellozia*, the most species-rich genus of the family. Using PacBio HiFi sequencing and Omni-C chromatin conformation data, we generated a haplotype-resolved genome assembly of *Vellozia tubiflora* with near telomere-to-telomere continuity. Hap1 and Hap2 span 423.2 Mb and 426.4 Mb, respectively, with both assemblies successfully anchored into nine chromosome-length scaffolds. The former presents a scaffold N50 of 49.8 Mb and 96.5% BUSCO completeness, while the latter reaches 50.0 Mb and 96.8%. Genome annotation predicted 23,352 and 23,549 protein-coding genes in Hap1 and Hap2, respectively. This resource is anticipated to foster new insights for comparative and evolutionary studies of vegetative desiccation tolerance across angiosperms and expands the genomic representation of plants from the Brazilian *campos rupestres*, a threatened biodiversity hotspot characterized by extreme environmental conditions.

**Significance:** The first haplotype-resolved genome of *V. tubiflora* provides a foundational resource for investigating the evolution and molecular basis of vegetative desiccation tolerance in the angiosperm lineage with the greatest number of resurrection species. As the most widely distributed species in Velloziaceae, *V. tubiflora* also represents a valuable model for comparative studies of genome evolution, structural variation, and adaptation across environmentally heterogeneous landscapes, while expanding the genomic representation of the Brazilian *campos rupestres*, a threatened ecosystem recognized for its exceptional biodiversity and high levels of endemism.

## Introduction

Vegetative desiccation tolerance is a distinct strategy that very few plant species exhibit to cope with long and/or severe drought episodes, characterized by their ability to survive the loss of up to 95% of their cellular water and fully resume metabolic and photosynthetic activities upon rehydration (Oliver, et al. 2020; Porembski and Barthlott 2000). While desiccation tolerance is widespread in orthodox seeds, its occurrence in vegetative tissues is restricted to a relatively small number of species, particularly among angiosperms, ∼ 600 in this > 300,000-species clade (Marks, et al. 2024; Porembski and Barthlott 2000). Because of its unique nature, this strategy has attracted increasing attention over the past decades as a model to investigate the molecular, physiological, and evolutionary mechanisms that enable plants to endure severe water deficit (Marks, et al. 2025).

In recent years, advances in long-read sequencing technologies have enabled the generation of high-quality reference genomes for several angiosperm species exhibiting vegetative desiccation tolerance (Gao, et al. 2024; Marks, et al. 2026b; Marks, et al. 2024). Among the monocots, genomes of desiccation-tolerant species are largely concentrated within Poales, particularly Poaceae (Gao, et al. 2024). Increasing genomic representation across monocot lineages will help reveal which features of vegetative desiccation tolerance are broadly conserved and which have evolved or diversified within particular lineages. These genomic resources have substantially expanded our understanding of the genetic architecture and evolutionary basis of plant adaptation to drought stress. However, taxonomic and geographic biases persist, and some of the most species-rich and ecologically unique desiccation-tolerant species still lack genome representation, limiting our ability to comprehend the mechanisms underlying desiccation tolerance across evolutionary lineages (Marks, et al. 2026a; Marks, et al. 2021).

The family Velloziaceae is, among angiosperms, the lineage with the largest number of vegetative desiccation-tolerant species (Alcântara, et al. 2015; Porembski and Barthlott 2000). It comprises five genera, *Acanthochlamys, Barbacenia, Barbaceniopsis, Vellozia*, and *Xerophyta* (Alcântara, et al. 2018). However, to date, available genome assemblies are restricted to the *Acanthochlamys* and *Xerophyta* genera (Costa, et al. 2017; Gao, et al. 2021; Xu, et al. 2022), leaving the most species-rich genera in the family underrepresented. Notably, *Vellozia*, the largest genus with approximately 128 described species (Alcântara, et al. 2018; Magri, et al. 2025), ∼ 70% displaying desiccation-tolerance traits (Larocca, et al. 2022), has remained entirely unexplored at the genomic level.

Both *Vellozia* and *Barbacenia* genera have their centers of diversity in the Brazilian *campos rupestres*, a threatened (Bugado, et al. 2025) montane ecosystem primarily distributed along the Espinhaço Range and Serra da Canastra in central-eastern Brazil presenting exceptional plant diversity and high levels of endemism (Alcântara, et al. 2018; Magri, et al. 2025). *Campo rupestre* vegetation typically thrives in oligotrophic environments characterized by limiting edaphic conditions such as sandy and rocky, highly draining, acidic and nutrient-poor substrates (especially in regard to phosphorus), droughts, and large daily thermal amplitudes and rates of evapotranspiration and radiance (Alcântara, et al. 2018; Oliveira, et al. 2015). Together, these conditions shaped a flora that is exceptionally rich in stress-adapted species (Zappi, et al. 2017).

Despite its ecological and evolutionary significance, this ecosystem remains underrepresented in plant genomics.

Here, we present the reference genome of the dissection-tolerant species *V. tubiflora* (Larocca, et al. 2022; Pinto, et al. 2025), to our knowledge a first for the genus. It is also the first haplotype-resolved assembly within the Velloziaceae family and achieves near telomere-to-telomere continuity. This level of completeness enables detailed analyses of genome structure, allelic variation, and repetitive landscape, providing new opportunities to investigate the evolutionary origins, conservation, and diversification of the molecular mechanisms underlying vegetative desiccation tolerance and adaptation to drought. In addition, while most species in the family are highly endemic (Magri, et al. 2025), *V. tubiflora* exhibits the broadest geographic distribution in the family, occurring from southeastern Brazil to Panama, across diverse altitudes, soils, and habitats (Mello-Silva 2010). This wide occurrence suggests a highly adaptable genetic background, making *V. tubiflora* an informative model for studying evolutionary diversification and molecular variation associated with desiccation tolerance.

## Results and Discussion

### Genome Size Estimation and Sequencing Resources for Genome Assembly

Using flow cytometry and *Arabidopsis thaliana* Col-0 as the internal standard, we estimated the haploid genome size of *V. tubiflora* at approximately 464 ± 1 Mb (Fig. S1) from three individuals accessed in a *campo rupestre* site in the Serra da Canastra National Park (Figs. 1a-c). Sequencing data was generated from a single individual. GenomeScope 2.0 (Ranallo-Benavidez, et al. 2020) k-mer based analysis estimated a smaller genome size of 402.76 Mb (Fig. S2), with a sequencing error rate of 0.255% and heterozygosity of 0.86%. The k-mer spectrum exhibited a bimodal distribution, with two major peaks at approximately 22x and 44x coverage, corresponding to heterozygous and homozygous k-mers, respectively (Fig. S2). The difference between the flow cytometry and k-mer-based genome size estimates may reflect underestimation by k-mer-based approaches in genomes with high repetitive content, particularly when repetitive k-mers are filtered or poorly represented in the fitted model (Li, et al. 2025). To generate a high-quality reference genome, we used approximately 21 Gb of PacBio HiFi reads, corresponding to ∼45x genome coverage based on the genome size estimated by flow cytometry. The HiFi dataset comprised 1.30 million reads, with a mean read length of 15,888 bp, an N50 read length of 16,180 bp, and read lengths ranging from 45 to 45,488 bp. For chromosome-scale scaffolding, three Omni-C libraries generated approximately 64 Gb of sequence data, comprising 212,852,036 read pairs.

**Figure 1.**
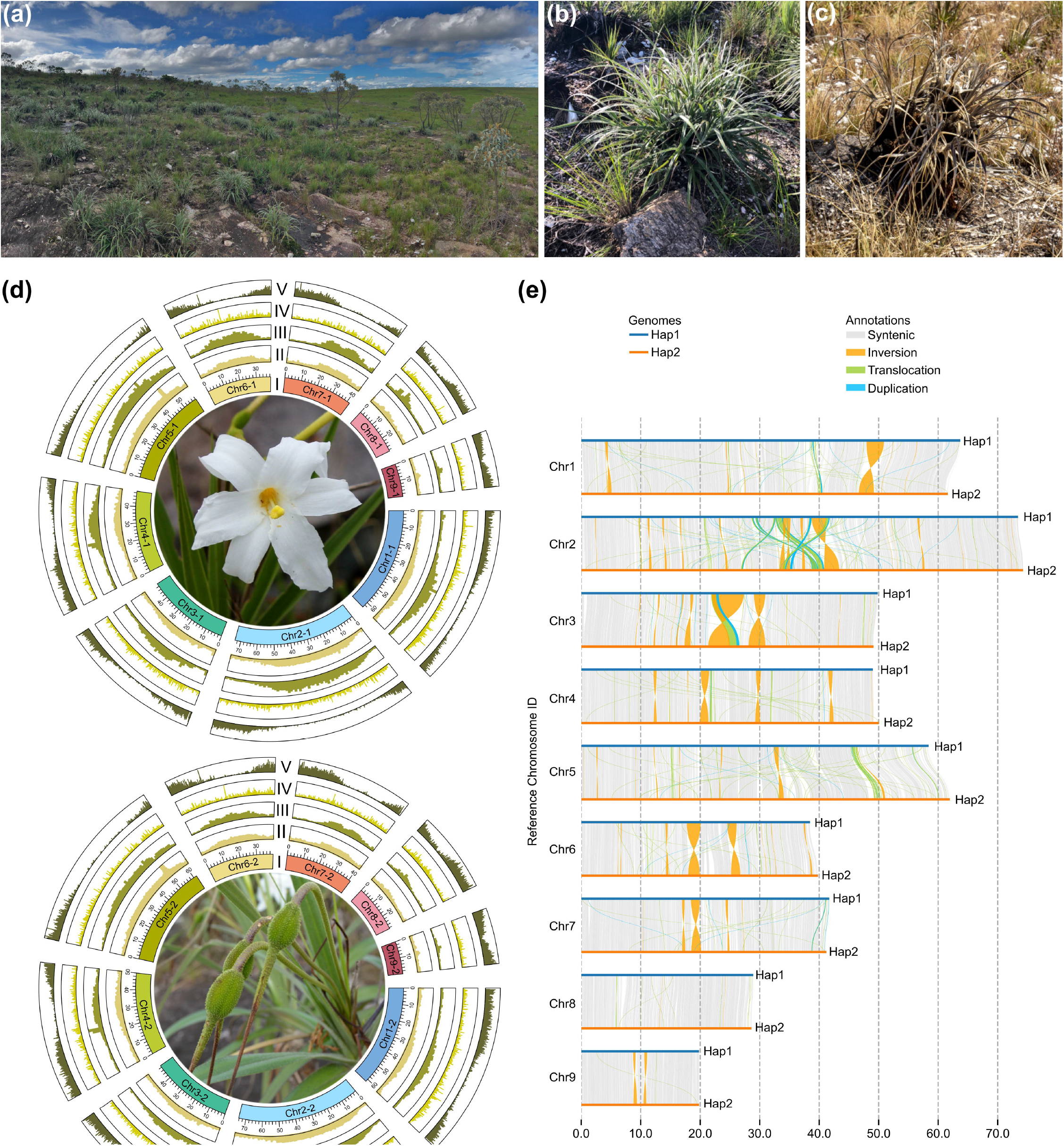
*Vellozia tubiflora* and its genome. a) *Campo rupestre* site at Serra da Canastra National Park accessed in this study (left; photograph by Isabel R. Gerhardt); hydrated *V. tubiflora* individual in the rainy season (center; photograph by Isabel R. Gerhardt); and desiccated individual in the dry season (right; photograph by Rafael S. C. de Souza). b) Circos plots of Hap1 (top) and Hap2 (bottom) were generated with TBtools-II. Features are represented by tracks labeled with Roman numerals: I, chromosomes (length expressed in Mb); II, GC content; III, Ty3-Gypsy and IV, Ty1-Copia LTR retrotransposon densities; V, gene density. Flower and immature fruit photographs adapted from original versions authored by, respectively, Mauricio Mercadante (https://www.inaturalist.org/observations/175940281; CC BY-NC-SA 4.0) and Pablo Preliasco (https://www.inaturalist.org/photos/63826408; CC BY-NC 4.0). c) Synteny and structural variation between the two *V. tubiflora* genome haplotypes. Chromosomes 1-9 of Hap1 and Hap2 were aligned, and genomic regions were classified as syntenic, inverted, translocated, duplicated (as indicated by chromatic colors) and haplotype-specific (white). Lengths are expressed in Mb.

### Genome Assembly

We generated a chromosome-level, haplotype-resolved *V. tubiflora* genome assembly with near telomere-to-telomere contiguity (Figs. 1d,e, S3 and S4). Detailed assembly statistics are reported in Table 1. Briefly, Hap1 consists of nine scaffolds spanning 423.2 Mb with contig N50 of 1.48 Mb, scaffold N50 of 49.8 Mb, longest contig of 7.56 Mb, and longest scaffold of 73.4 Mb. GC content is 40.67%. Hap2 comprises nine scaffolds spanning 426.5 Mb with contig N50 of 1.52 Mb, scaffold N50 of 50.0 Mb, longest contig of 8.7 Mb, and longest scaffold of 74.3 Mb. GC content for Hap2 is 40.74%. The nine chromosome-length scaffolds are consistent with the basic chromosome number reported for the genus *Vellozia* (Braz, et al. 2025). These chromosome-scale assemblies represent approximately 92% of the genome size estimated by flow cytometry. An additional 43.1 Mb of sequence remained unplaced and could not be confidently assigned to a chromosome or phased to either haplotype. Telomeric motifs (AAACCCT)n were identified at 26 of the 36 chromosome ends, with at least one telomeric end in each chromosome (Table S1 and Fig. S4), supporting the classification of both haplotype assemblies as near telomere-to-telomere. Genome completeness assessment using BUSCO (Manni, et al. 2021) showed 96.5% and 96.8% complete BUSCOs (embryophyta_odb10 database) for Hap1 and Hap2, respectively. Most complete BUSCOs were single-copy (94.1% in both haplotypes), with only 2.4% and 2.7% duplicated BUSCOs in Hap1 and Hap2. The assemblies exhibit Merqury consensus quality values (QV) of 66.46 and 66.28 for Hap1 and Hap2, respectively, with k-mer completeness of 83.97% and 83.96%, respectively, and 96.85% for the combined assembly. LTR Assembly Index (LAI) values of 23.47 and 23.54, respectively, further support the high accuracy and contiguity of the assemblies across repetitive regions.

**Table 1.**
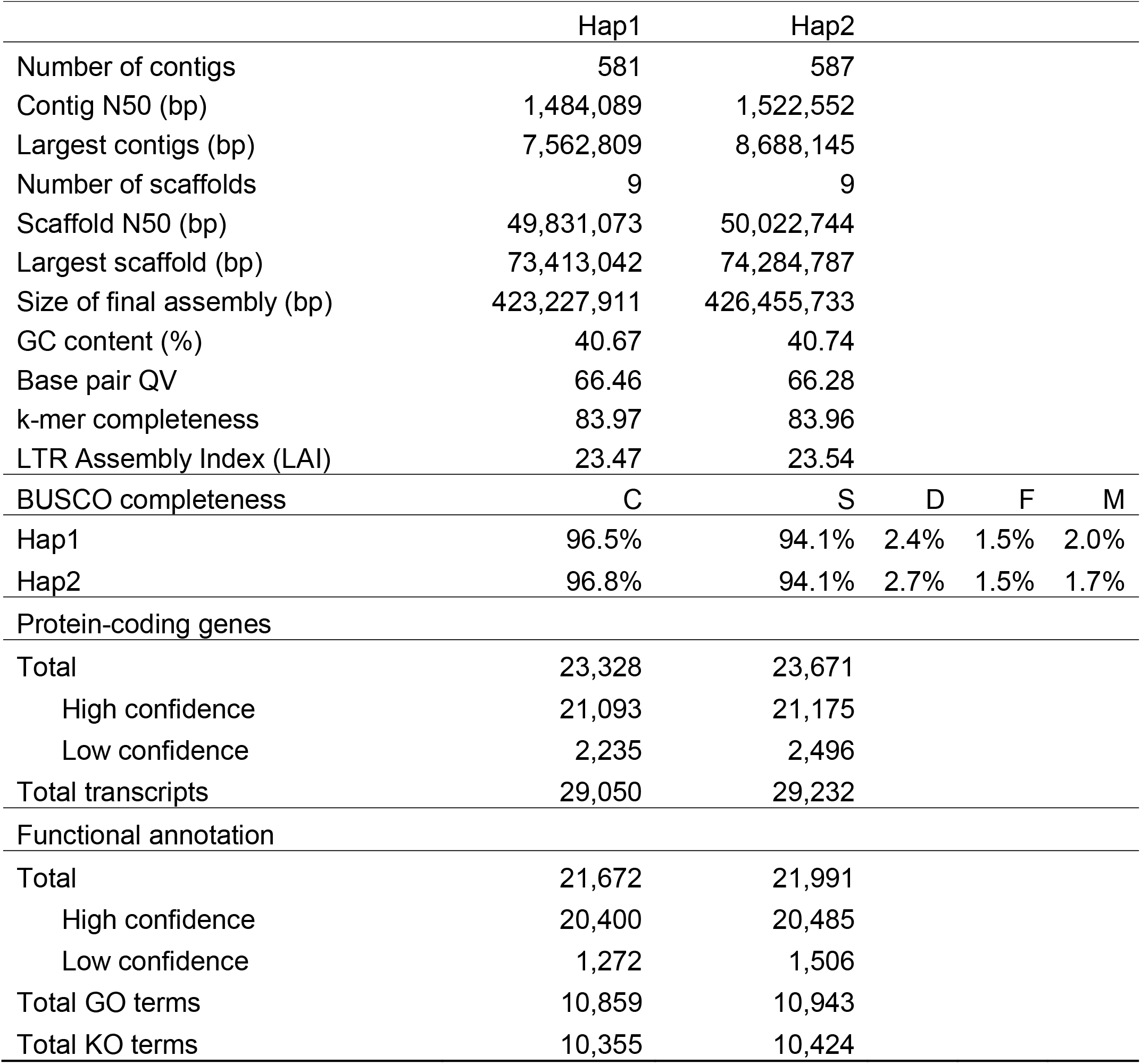
Summary of the *V. tubiflora* genome assembly, annotation, and quality assessment.

|  | Hap1 | Hap2 |  |  |  |
| --- | --- | --- | --- | --- | --- |
| Number of contigs | 581 | 587 |  |  |  |
| Contig N50 (bp) | 1,484,089 | 1,522,552 |  |  |  |
| Largest contigs (bp) | 7,562,809 | 8,688,145 |  |  |  |
| Number of scaffolds | 9 | 9 |  |  |  |
| Scaffold N50 (bp) | 49,831,073 | 50,022,744 |  |  |  |
| Largest scaffold (bp) | 73,413,042 | 74,284,787 |  |  |  |
| Size of final assembly (bp) | 423,227,911 | 426,455,733 |  |  |  |
| GC content (%) | 40.67 | 40.74 |  |  |  |
| Base pair QV | 66.46 | 66.28 |  |  |  |
| k-mer completeness | 83.97 | 83.96 |  |  |  |
| LTR Assembly Index (LAI) | 23.47 | 23.54 |  |  |  |
| BUSCO completeness | C | S | D | F | M |
| Hap1 | 96.5% | 94.1% | 2.4% | 1.5% | 2.0% |
| Hap2 | 96.8% | 94.1% | 2.7% | 1.5% | 1.7% |
| Protein-coding genes |  |  |  |  |  |
| Total | 23,328 | 23,671 |  |  |  |
| High confidence | 21,093 | 21,175 |  |  |  |
| Low confidence | 2,235 | 2,496 |  |  |  |
| Total transcripts | 29,050 | 29,232 |  |  |  |
| Functional annotation |  |  |  |  |  |
| Total | 21,672 | 21,991 |  |  |  |
| High confidence | 20,400 | 20,485 |  |  |  |
| Low confidence | 1,272 | 1,506 |  |  |  |
| Total GO terms | 10,859 | 10,943 |  |  |  |
| Total KO terms | 10,355 | 10,424 |  |  |  |

### Genome Annotation

The annotation of transposable elements (TEs) was conducted using the EDTA pipeline (Ou, et al. 2019a), and identified a repeat rich genome with repetitive sequences comprising 65.97% (279.2 Mb) and 66.27% (282.6 Mb) for Hap1 and Hap2, respectively (Table S2). In Hap1, long terminal repeats retrotransposons (LTR) were the predominant class, accounting for 50.11% of the assembly, with Ty3-Gypsy and Ty1-Copia elements representing 38.49% and 5.13% of the genome, respectively. In Hap2, LTR retrotransposons comprise 48.67% of the genome, with Ty3-Gypsy and Ty1-Copia accounting for 36.42% and 5.92% of the assembly, respectively. DNA transposons account for 8.65% and 8.90% of the genome assemblies for Hap1 and Hap2, respectively. These genomic features are represented across the nine chromosome-length pseudochromosomes in the *V. tubiflora* circos plot (Fig. 1d) generated with TBtools-II (Chen, et al. 2023).

Integration of Helixer predictions (Holst, et al. 2026a) and BRAKER4 (ETP mode; Gabriel et al., 2024) resulted in the final annotations of 23,328 and 23,671 protein-coding genes in Hap1 and Hap2, respectively. These genes were classified into high- and low-confidence according to transcriptomic and protein evidence (Table 1). The predicted proteomes showed 97.6% BUSCO completeness against the embryophyta_odb10 dataset in both haplotypes. Functional annotation using eggNOG-mapper resulted in orthology assignments for 92.9% genes in both haplotypes, of which 46.5% and 46.2% were associated to Gene Ontology (GO) terms, and 44.3 and 44.0% to KEGG Orthology (KO) terms (Table 1, Table S3 and S4).

### Structural Variation Between the Two *V. tubiflora* Haplotypes

To characterize structural variation between haplotypes, Hap1 and Hap2 were aligned using Nucmer, and syntenic relationships and structural rearrangements were identified with SyRI. A total of 11,113 syntenic regions were identified, spanning approximately 301.5 Mb in Hap1 and 302.7 Mb in Hap2 (Fig. 1e and Table S5), with 71.2% and 70.9% of the assembled sequences classified as syntenic in Hap1 and Hap2, respectively. The analysis also revealed 155 inversions (∼30 Mb) and 9,702 translocations (∼37 Mb) between the two assemblies. Large inversions were observed on chromosomes 1, 2, 3, 6 and 7 (Fig.1e). In addition, we identified 2,133 duplications in Hap1 (∼8.8 Mb) and 2,015 duplications in Hap2 (∼7.2 Mb) (Fig. 1e and Table S5). This analysis yielded a total of 2,245,742 single-nucleotide polymorphism (SNP) differences, 303,366 insertions (∼8.3 Mb), and 245,989 deletions (∼7.6 Mb) (Table S5). As the first haplotype-resolved genome in the Velloziaceae, this assembly provides a foundation for investigating the evolutionary and functional significance of intraspecific structural variation.

### Material and Methods Plant Materials

Three *V. tubiflora* individuals were accessed from a single population located in a *campo rupestre* site in Serra da Canastra National Park, in the municipality of São Roque de Minas in southern Minas Gerais state, Brazil. Coordinates and elevation (in m.a.s.l.) of this site are S 20º 13’50.8” W 46º 37’31.2”, 1,380.

### Measurement of Genome Size Through Flow Cytometry

Genome size was estimated by flow cytometry using *A. thaliana* Col-0 (2C = 314 Mb) (Bennett, et al. 2003) as an internal standard. Leaf tissue from the three accessed *V. tubiflora* individuals and *A. thaliana* was separately chopped as previously described (Dilkes, et al. 2002) in propidium iodide (PI) and RNase staining buffer (Ebihara, et al. 2005), filtered, incubated at 37°C for 15 min, and analyzed using a FACSCanto II flow cytometer (BD Biosciences) equipped with a 488 nm laser. Subsequently, each *V. tubiflora* homogenate was pooled with the *A. thaliana* homogenate, and fluorescence signals from 10,000 nuclei were acquired. The haploid (1C) genome size of *V. tubiflora* was calculated from the relative fluorescence intensity of sample and standard nuclei.

### DNA Extraction

High-molecular-weight genomic DNA (HMW gDNA) was extracted from young leaves of a single *V. tubiflora* individual using a modified cetyltrimethylammonium bromide (CTAB) protocol based on Doyle and Doyle (1987). The DNA size distribution and integrity were assessed using a Femto Pulse system (Agilent Technologies), while DNA concentration and purity were evaluated using a Qubit 2.0 fluorometer and a NanoDrop 2000c spectrophotometer, respectively (Thermo Fisher Scientific).

### HiFi Library Preparation and Sequencing

The PacBio HiFi library was prepared from HMW gDNA following the SMRTbell Express Template Prep Kit v2.0 protocol (PacBio). Genomic DNA was fragmented to approximately 15–20 kb using Diagenode’s Megaruptor 3 system (Diagenode) and concentrated with AMPure PB beads before library construction. SMRTbell preparation included removal of single-stranded overhangs, DNA damage and end repair, A-tailing, adapter ligation, and nuclease treatment. Following purification, the library was size selected using the BluePippin/PippinHT system (Sage Science), retaining fragments larger than approximately 9 kb. The resulting library, with an average fragment size of 15–20 kb, was sequenced at the UC Davis DNA Technologies Core on a PacBio Sequel II platform using one 8M SMRT Cell, Sequel II chemistry 2.0, and 30-h movies per cell.

### Omni-C Library Preparation and Sequencing

Three Omni-C libraries were prepared using the Dovetail Omni-CTM Kit (Dovetail Genomics) according to the manufacturers protocol. Briefly, chromatin was fixed in place with formaldehyde in the nucleus and then extracted. Fixed chromatin was digested with DNAse I, chromatin ends were repaired and ligated to a biotinylated bridge adapter followed by proximity ligation of adapter containing ends. After proximity ligation, crosslinks were reversed and the DNA purified. Purified DNA was treated to remove biotin that was not internal to ligated fragments. Sequencing libraries were generated using NEBNext Ultra enzymes and Illumina-compatible adapters. Biotin-containing fragments were isolated using streptavidin beads before PCR enrichment of each library. Libraries were sequenced on an Illumina HiSeqX platform to generate approximately 213 million 2 x 150 bp read pairs.

### RNA extraction

Root, stem, and leaf tissues were collected from three *V. tubiflora* individuals growing in a *campo rupestre* site in Serra da Canastra National Park, Minas Gerais, Brazil. Fresh tissues were immediately frozen in dry ice and stored at −80 °C for further RNA extraction. Total RNA was extracted independently from each tissue and individual using a CTAB-based protocol followed by purification with the RNeasy Plant Mini Kit (Qiagen). RNA concentration and quality were assessed prior to library preparation, and only samples with an RNA integrity number (RIN) greater than 7 were used for sequencing.

### RNA-seq library preparation and sequencing

A total of nine RNA-seq libraries were prepared, representing three biological replicates for each tissue (root, stem, and leaf). Stranded mRNA libraries were generated using the Illumina Stranded mRNA Prep, Ligation kit according to the manufacturer’s instructions. Libraries were sequenced on an Illumina NextSeq 2000 platform using a P3 200-cycle sequencing kit to generate 2 x 100bp paired-end reads.

### Nuclear Genome Assembly

HiFi reads were assembled using hifiasm v0.20.0 (Cheng, et al. 2021) with the HiC read integration parameters. The assembled contigs were then filtered for contamination using v0.5.4 of NCBI’s Foreign Contamination Screening – GX package (Astashyn, et al. 2024). The filtered contigs were subsequently scaffolded using the Omni-C data and HapHiC v1.0.6 (Zeng, et al. 2024) with manual curation using Juicebox v1.11.08 (Durand, et al. 2016).

### Genome Annotation

Initial transposable element annotation of the genome assembly was conducted using The Extensive de novo TE Annotator (EDTA) v1.9.6 (Ou, et al. 2019b). Protein-coding genes were predicted independently with Helixer v0.3.5 using the land_plant lineage parameter (Holst, et al. 2026b), and BRAKER4 v.0.5.0-beta (https://github.com/Gaius-Augustus/BRAKER4). BRAKER4 was run in ETP mode (Gabriel, et al. 2024), using RNA-seq reads from root, stem, and leaf tissues mapped to each haplotype assembly with HISAT2 v.2.2.1, and Viridiplantae protein evidence from OrthoDB (Tegenfeldt, et al. 2025). Genes exclusively predicted by Helixer were identified using gffcompare v.0.12.10 (Pertea and Pertea 2020) from class “u”.

BRAKER4 and Helixer-specific predictions were classified into high-(HC) and low-confidence (LC) sets based on transcriptomic and protein evidence. Predicted proteins were searched with DIAMOND v.2.1.10 (Buchfink, et al. 2021) against the plant subset of Swiss-Prot, TrEMBL and the whole PTREP (Release 25; https://trep-db.uzh.ch/) using an E-value threshold of 1 e-10. Swiss-Prot and TrEMBL matches with ≥ 80% query and subject coverage were considered strong protein support, while PTREP matches covering ≥ 70% of the query were considered evidence of a TE-related protein.

For BRAKER4 predictions, transcript-level structural support was evaluated using the evidence reported by BRAKER4. Multi-exon transcripts were considered structurally HC when ≥ 50% of introns were supported by RNA-seq or protein evidence, or, in the absence of intron support, when exon support was complete. Single-exon transcripts required complete RNA-seq or protein support. Genes with at least one supported transcript were classified as HC, while unsupported genes were retained as HC when supported by strong protein homology. Genes with strong PTREP matches were classified as LC.

For Helixer-specific models, RNA-seq support was assessed from exonic coverage and splice-junction support. Splice junctions were extracted with RegTools v.1.0.0 (Cotto, et al. 2023) and converted to intron coordinates, which were exactly matched by position and strand to predicted introns using BEDTools v.2.31.1 (Quinlan and Hall 2010). Exonic support required ≥ 50% coverage of total predicted exonic bases and ≥ 10 exon-level alignment overlaps. Models were classified as HC when they had (i) a strong SwissProt match, (ii) a strong TrEMBL match combined with exonic or exact splice-junction support, or (iii) exonic support combined with at least one exact splice-junction match. Models with strong PTREP matches were classified as LC unless supported by Swiss-Prot. All remaining models were assigned to the LC set. HC and LC annotations from both approaches were merged separately, and models with incomplete coding sequences were removed using AGAT v.1.7.0 (Dainat, et al. 2026). Detailed classification criteria and scripts are available at [https://github.com/GCCRC-dev/annotation-BRAKER-Helixer].

Functional annotation was performed on the longest protein isoform of each gene using eggNOG-mapper v.2.1.12 (Cantalapiedra, et al. 2021) with the eggNOG database v.5.0.2.

### Telomere Repeats Annotation

Telomeric repeat arrays were identified using the QuarTeT toolkit with the -c plant option to screen the assembled genome for the plant telomeric repeat (AAACCCT)n (Lin, et al. 2023).

### Structural Variation Between Haplotypes

To identify structural rearrangements between the two haplotype assemblies of *V. tubiflora*, whole-genome alignments were generated using NUCmer, implemented in MUMmer4 (v 3.1) (Kurtz, et al. 2004; Marçais, et al. 2018), with the parameters --mum -l 50 -b 500 -c 200. The resulting delta alignments were filtered using delta-filter to retain alignments of at least 200 bp and with a minimum sequence identity of 95% (-l 200 -i 95 -r -q). Filtered alignments were converted to tab-delimited coordinates using show-coords, and structural rearrangements were identified using SyRI (v1.7.1) with default parameters (Goel, et al. 2019). The resulting syntenic regions and structural rearrangements were visualized using plotsr (v 1.1.5) (Goel and Schneeberger 2022).

## Supplementary Material

Supplementary material is available online.

## Acknowledgments

We thank the Life Sciences Core Facility (LaCTAD) at the Universidade Estadual de Campinas for flow cytometric analysis.

## Author Contributions

RAD and IRG conceived and designed the study. MFC, RAD, and IRG collected and identified plant material for genomic analyses. MFC, OHN, and RS performed laboratory work and generated sequencing data. LECC, ME, DMRP, and SR performed preliminary genome assembly. NAA and TPM generated the final genome assemblies and annotations, with MPM contributing to the latter. FANA and MPM performed statistical analyses. FANA, NAA, MPM, TPM, RAD, and IRG wrote the initial manuscript draft. HAL and PA provided infrastructure and financial support. All authors reviewed, edited, and approved the final manuscript.

## Funding

This work was supported by grants 2016/23218-0 from Fundação de Amparo à Pesquisa do Estado de São Paulo (FAPESP) and Seed Grant for International Activities from Global Affairs and the College/School of Biological Sciences from University of California, Davis. FANA (2024/11068-0 and 2026/00072-1), LECC (2018/19099-0), MFC (2022/049293) and MPM (2026/15267-2) received fellowships from FAPESP.

## Conflict of Interest

The authors declare no conflict of interest.

## Data Availability

Data generated for this study are available under NCBI BioProject PRJNA1530602.

## Legends

**Figure S1**. Estimation of *V. tubiflora* genome sizes through flow cytometry. Leaf homogenates containing nuclei were prepared from *V. tubiflora* individuals and the standard *A. thaliana* Col-0, treated with ribonuclease and stained with PI. The homogenates of *V. tubiflora* were individually pooled with that of *A. thaliana* and examined by flow cytometry. a) Uniparametric distributions of fluorescence emission in linear scale of a *V. tubiflora* individual and *A. thaliana* standard. *At* and *Vt*, peaks corresponding to *A. thaliana* and *V. tubiflora* nuclei, respectively; C, haploid nuclear DNA content. *A. thaliana* leaf homogenate yields both 2C and endoreduplicated (4C and 8C) nuclei; conversely, *V. tubiflora* leaf homogenate yields a single peak corresponding to 2C nuclei with DNA content sized between *A. thaliana* 4C (628 Mb) and 8C (1,256 Mb) peaks. b) Samples, standards, PI florescence values and calculated 2C and C genome sizes (expressed in Mb) of the three *V. tubiflora* individuals using the 2C genome content of the *A. thaliana* Col-0 standard.

**Figure S2**. K-mer profile (k = 51) spectral analysis to estimate genome size of *V. tubiflora*. The two peaks represent two distinct k-mer frequency distributions: The first peak (lower frequency) corresponds to heterozygous k-mers, suggesting variation between homologous chromosomes. The second peak (higher frequency) represents homozygous k-mers, which are present in both copies of the genome.

**Figure S3**. Omni-C contact heatmap of the haplotype-resolved *V. tubiflora* genome assembly, displaying chromatin interaction frequencies across the scaffolds of Hap1 and Hap2.

**Figure S4**. Gene density and telomeric repeat distribution in the *V. tubiflora* genome. Gene density across the nine chromosome-length scaffolds of (a) Hap1 and (b) Hap2 is represented by the color gradient from low to high density. Black arrowheads indicate chromosome ends containing telomeric repeat arrays (AAACCCT)n.

## Notes

### Competing Interest Statement

The authors have declared no competing interest.

## Literature Cited

Alcântara S, et al. 2015. Carbon assimilation and habitat segregation in resurrection plants: a comparison between desiccation- and non-desiccation-tolerant species of Neotropical Velloziaceae (Pandanales). Functional Ecology 29: 1499–1512. doi: 10.1111/1365-2435.12462

Alcântara S, Ree RH, Mello-Silva R 2018. Accelerated diversification and functional trait evolution in Velloziaceae reveal new insights into the origins of the campos rupestres’ exceptional floristic richness. Ann Bot 122: 165–180. doi: 10.1093/aob/mcy063

Astashyn A, et al. 2024. Rapid and sensitive detection of genome contamination at scale with FCS-GX. Genome Biology 25: 60. doi: 10.1186/s13059-024-03198-7

Bennett MD, Leitch IJ, Price HJ, Johnston JS 2003. Comparisons with Caenorhabditis (approximately 100 Mb) and Drosophila (approximately 175 Mb) using flow cytometry show genome size in Arabidopsis to be approximately 157 Mb and thus approximately 25% larger than the Arabidopsis genome initiative estimate of approximately 125 Mb. Ann Bot 91: 547–557. doi: 10.1093/aob/mcg057

Braz GT, et al. 2025. Revisiting the cytogenetics of Vellozia Vand.: immunolocalization of KNL1 elucidates the chromosome number for the genus. Brazilian Journal of Botany 48: 6. doi: 10.1007/s40415-024-01064-1

Buchfink B, Reuter K, Drost HG 2021. Sensitive protein alignments at tree-of-life scale using DIAMOND. Nat Methods 18: 366–368. doi: 10.1038/s41592-021-01101-x

Bugado RE, Shrestha N, Magri RA, Prado J, Lopes JC 2025. Vanishing ecosystems: The looming threat of climate change on an iconic genus Vellozia in the Brazilian campos rupestres. Global Ecology and Conservation 58: e03439. doi: 10.1016/j.gecco.2025.e03439

Cantalapiedra CP, Hernandez-Plaza A, Letunic I, Bork P, Huerta-Cepas J 2021. eggNOG-mapper v2: Functional Annotation, Orthology Assignments, and Domain Prediction at the Metagenomic Scale. Mol Biol Evol 38: 5825–5829. doi: 10.1093/molbev/msab293

Chen C, et al. 2023. TBtools-II: A ‘‘one for all, all for one’’ bioinformatics platform for biological big-data mining. Molecular Plant 16: 1733–1742. doi: 10.1016/j.molp.2023.09.010

Cheng H, Concepcion GT, Feng X, Zhang H, Li H 2021. Haplotype-resolved de novo assembly using phased assembly graphs with hifiasm. Nat Methods 18: 170–175. doi: 10.1038/s41592-020-01056-5

Costa MCD, et al. 2017. A footprint of desiccation tolerance in the genome of Xerophyta viscosa. Nature Plants 3: 17038. doi: 10.1038/nplants.2017.38

Cotto KC, et al. 2023. Integrated analysis of genomic and transcriptomic data for the discovery of splice-associated variants in cancer. Nat Commun 14: 1589. doi: 10.1038/s41467-023-37266-6

Dainat J, et al. 2026. NBISweden/AGAT: AGAT v1.7.0 (Version v1.7.0) [Computer software]. Version 1.7.0: Zenodo.

Dilkes BP, Dante RA, Coelho C, Larkins BA 2002. Genetic analyses of endoreduplication in Zea mays endosperm: evidence of sporophytic and zygotic maternal control. Genetics 160: 1163–1177. doi: 10.1093/genetics/160.3.1163

Doyle JJ, Doyle JL 1987. A Rapid DNA Isolation Procedure for Small Quantities of Fresh Leaf Tissue. Phytochemical Bulletin 19: 11–15.

Durand NC, et al. 2016. Juicebox Provides a Visualization System for Hi-C Contact Maps with Unlimited Zoom. Cell Systems 3: 99–101. doi: 10.1016/j.cels.2015.07.012

Ebihara A, et al. 2005. Nuclear DNA, chloroplast DNA, and ploidy analysis clarified biological complexity of the Vandenboschia radicans complex (Hymenophyllaceae) in Japan and adjacent areas. American Journal of Botany 92: 1535–1547. doi: 10.3732/ajb.92.9.1535

Gabriel L, et al. 2024. BRAKER3: Fully automated genome annotation using RNA-seq and protein evidence with GeneMark-ETP, AUGUSTUS, and TSEBRA. Genome Res 34: 769–777. doi: 10.1101/gr.278090.123

Gao B, et al. 2024. Drying without dying: A genome database for desiccation-tolerant plants and evolution of desiccation tolerance. Plant Physiol 194: 2249–2262. doi: 10.1093/plphys/kiad672

Gao ZY, Li ZH, Lin DL, Jin XH 2021. Chromosome-Scale Genome Assembly of the Resurrection Plant Acanthochlamys bracteata (Velloziaceae). Genome Biol Evol 13. doi: 10.1093/gbe/evab147

Goel M, Schneeberger K 2022. plotsr: visualizing structural similarities and rearrangements between multiple genomes. Bioinformatics 38: 2922–2926. doi: 10.1093/bioinformatics/btac196

Goel M, Sun H, Jiao WB, Schneeberger K 2019. SyRI: finding genomic rearrangements and local sequence differences from whole-genome assemblies. Genome Bio120: 277. doi: 10.1186/s13059-019-1911-0

Holst F, et al. 2026a. Helixer: ab initio prediction of primary eukaryotic gene models combining deep learning and a hidden Markov model. Nat Methods 23: 732–739. doi: 10.1038/s41592-025-02939-1

Holst F, et al. 2026b. Helixer: ab initio prediction of primary eukaryotic gene models combining deep learning and a hidden Markov model. Nat Methods 23: 732–739. doi: 10.1038/s41592-025-02939-1

Kurtz S, et al. 2004. Versatile and open software for comparing large genomes. Genome Biology 5: R12. doi: 10.1186/gb-2004-5-2-r12

Larocca PDF, Saldanha Mancio J, Padilha P, Mello-Silva R, Alcântara S 2022. Recent divergence in functional traits affects rates of speciation in the Neotropical Velloziaceae (Pandanales). Botanical Journal of the Linnean Society 199: 144–172. doi: 10.1093/botlinnean/boab102

Li M, et al. 2025. Is it time to abandon the flow cytometry in estimations of genome size when the K-mer analysis is available? The case of Engelhardia species. Genomics Communications 2: e013. doi: 10.48130/gcomm-0025-0014

Lin Y, et al. 2023. quarTeT: a telomere-to-telomere toolkit for gap-free genome assembly and centromeric repeat identification. Hortic Res 10: uhad127. doi: 10.1093/hr/uhad127

Magri RA, et al. 2025. Historical biogeography of Vellozia (Velloziaceae) reveals range expansion in South American mountaintops after climatic cooling events and increased diversification rates after the occupation of Southern Espinhaço Province. Botanical Journal of the Linnean Society 207: 115–127. doi: 10.1093/botlinnean/boae072

Manni M, Berkeley MR, Seppey M, Simão FA, Zdobnov EM 2021. BUSCO Update: Novel and Streamlined Workflows along with Broader and Deeper Phylogenetic Coverage for Scoring of Eukaryotic, Prokaryotic, and Viral Genomes. Mol Biol Evol 38: 4647–4654. doi: 10.1093/molbev/msab199

Marçais G, et al. 2018. MUMmer4: A fast and versatile genome alignment system. PLOS Computational Biology 14: e1005944. doi: 10.1371/journal.pcbi.1005944

Marks RA, Abrahams RS, Ekwealor JTB 2026a. Shaped by context: Evolutionary trajectories of desiccation tolerance in land plants. Am J Bot 113: e70180. doi: 10.1002/ajb2.70180

Marks RA, et al. 2025. Life on the dry side: a roadmap to understanding desiccation tolerance and accelerating translational applications. Nat Commun 16: 3284. doi: 10.1038/s41467-025-58656-y

Marks RA, Farrant JM, Nicholas McLetchie D, VanBuren R 2021. Unexplored dimensions of variability in vegetative desiccation tolerance. American Journal of Botany 108: 346–358. doi: 10.1002/ajb2.1588

Marks RA, et al. 2026b. The architecture of resilience: a genome assembly of Myrothamnus flabellifolia sheds light on desiccation tolerance and sex determination. New Phytol 249: 1063–1084. doi: 10.1111/nph.70700

Marks RA, Van Der Pas L, Schuster J, Gilman IS, VanBuren R 2024. Convergent evolution of desiccation tolerance in grasses. Nature Plants 10: 1112–1125. doi: 10.1038/s41477-024-01729-5

Mello-Silva Rd 2010. Circumscribing Vellozia hirsuta and V. tubiflora (Velloziaceae). Hoehnea 37: 617–646. doi: 10.1590/S2236-89062010000300009

Oliveira RS, et al. 2015. Mineral nutrition of campos rupestres plant species on contrasting nutrient-impoverished soil types. New Phytologist 205: 1183–1194. doi: 10.1111/nph.13175

Oliver MJ, et al. 2020. Desiccation Tolerance: Avoiding Cellular Damage During Drying and Rehydration. Annual Review of Plant Biology 71: 435–460. doi: 10.1146/annurev-arplant-071219-105542

Ou S, et al. 2019a. Benchmarking transposable element annotation methods for creation of a streamlined, comprehensive pipeline. Genome Biol 20: 275. doi: 10.1186/s13059-019-1905-y

Ou S, et al. 2019b. Benchmarking transposable element annotation methods for creation of a streamlined, comprehensive pipeline. Genome Biology 20: 275. doi: 10.1186/s13059-019-1905-y

Pertea G, Pertea M 2020. GFF Utilities: GffRead and GffCompare. F1000Res 9. doi: 10.12688/f1000research.23297.2

Pinto OHB, et al. 2025. Seasonal bacterial profiles of Vellozia with distinct drought adaptations in the megadiverse campos rupestres. Scientific Data 12: 636. doi: 10.1038/s41597-025-04984-z

Porembski S, Barthlott W 2000. Granitic and gneissic outcrops (inselbergs) as centers of diversity for desiccation-tolerant vascular plants. Plant Ecology 151: 19–28. doi: 10.1023/A:1026565817218

Quinlan AR, Hall IM 2010. BEDTools: a flexible suite of utilities for comparing genomic features. Bioinformatics 26: 841–842. doi: 10.1093/bioinformatics/btq033

Ranallo-Benavidez TR, Jaron KS, Schatz MC 2020. GenomeScope 2.0 and Smudgeplot for reference-free profiling of polyploid genomes. Nat Commun 11: 1432. doi: 10.1038/s41467-020-14998-3

Tegenfeldt F, et al. 2025. OrthoDB and BUSCO update: annotation of orthologs with wider sampling of genomes. Nucleic Acids Res 53: D516–D522. doi: 10.1093/nar/gkae987

Xu B, et al. 2022. Chromosome-level de novo genome assembly and whole-genome resequencing of the threatened species Acanthochlamys bracteata (Velloziaceae) provide insights into alpine plant divergence in a biodiversity hotspot. Mol Ecol Resour 22: 1582–1595. doi: 10.1111/1755-0998.13562

Zappi DC, Moro MF, Meagher TR, Nic Lughadha E 2017. Plant Biodiversity Drivers in Brazilian Campos Rupestres: Insights from Phylogenetic Structure. Front Plant Sci 8: 2141–2141. doi: 10.3389/fpls.2017.02141

Zeng X, et al. 2024. Chromosome-level scaffolding of haplotype-resolved assemblies using Hi-C data without reference genomes. Nature Plants 10: 1184–1200. doi: 10.1038/s41477-024-01755-3

