## Supplementary figures and images for "A haplotype-resolved, near telomere-to-telomere genome assembly of the desiccation-tolerant plant *Vellozia tubiflora*"

### Supplemental Figure 1

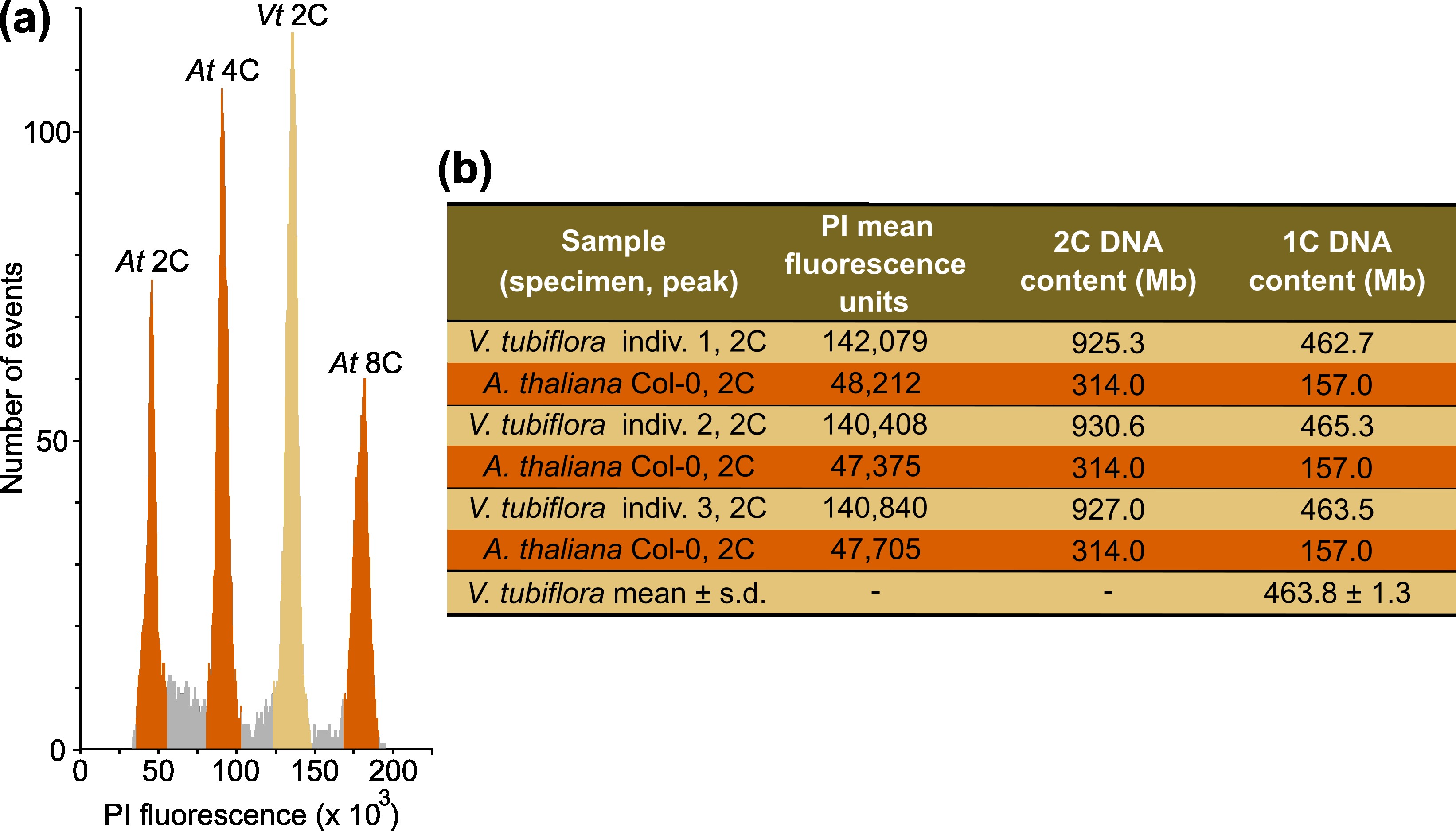

### Supplemental Figure 2

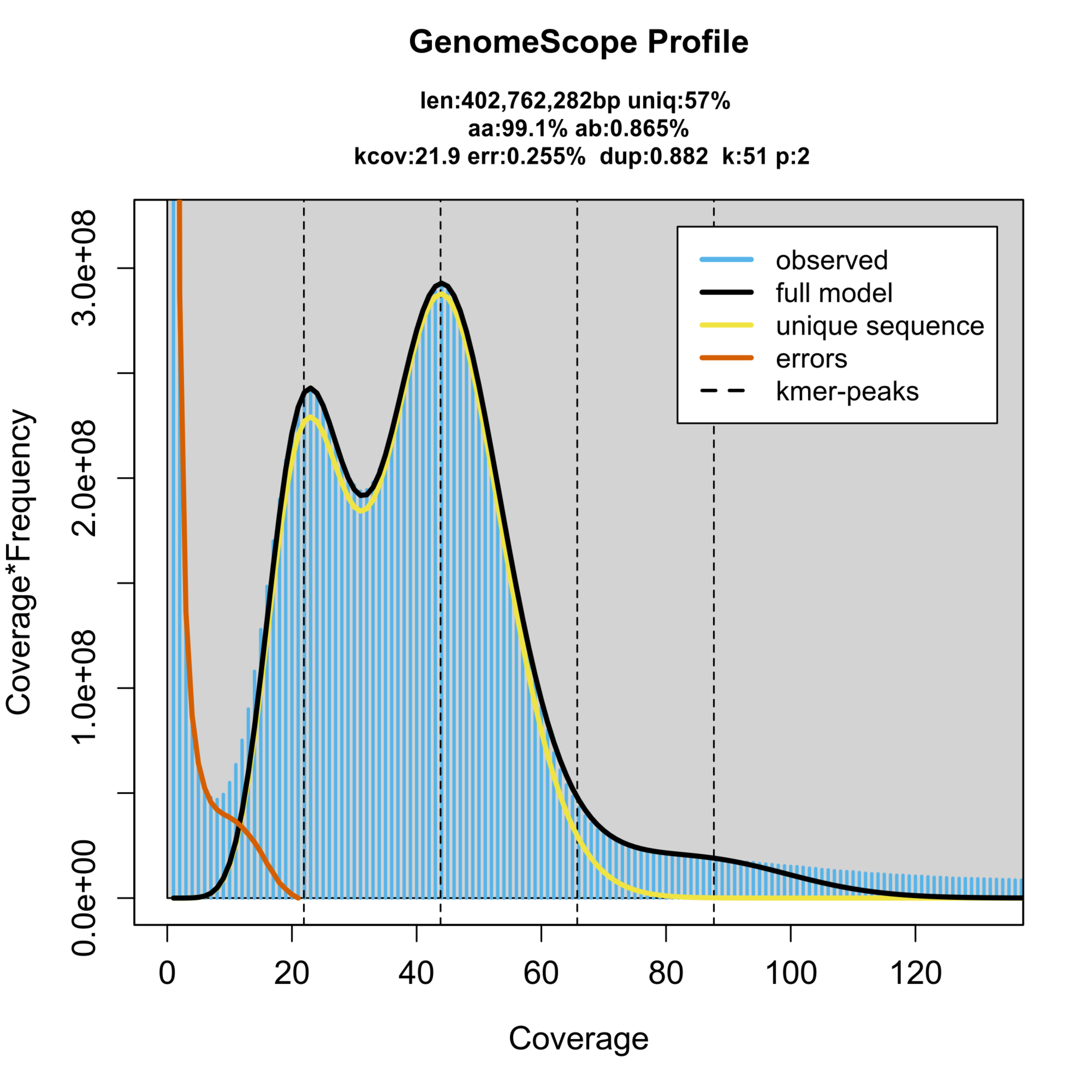

### Supplemental Figure 3

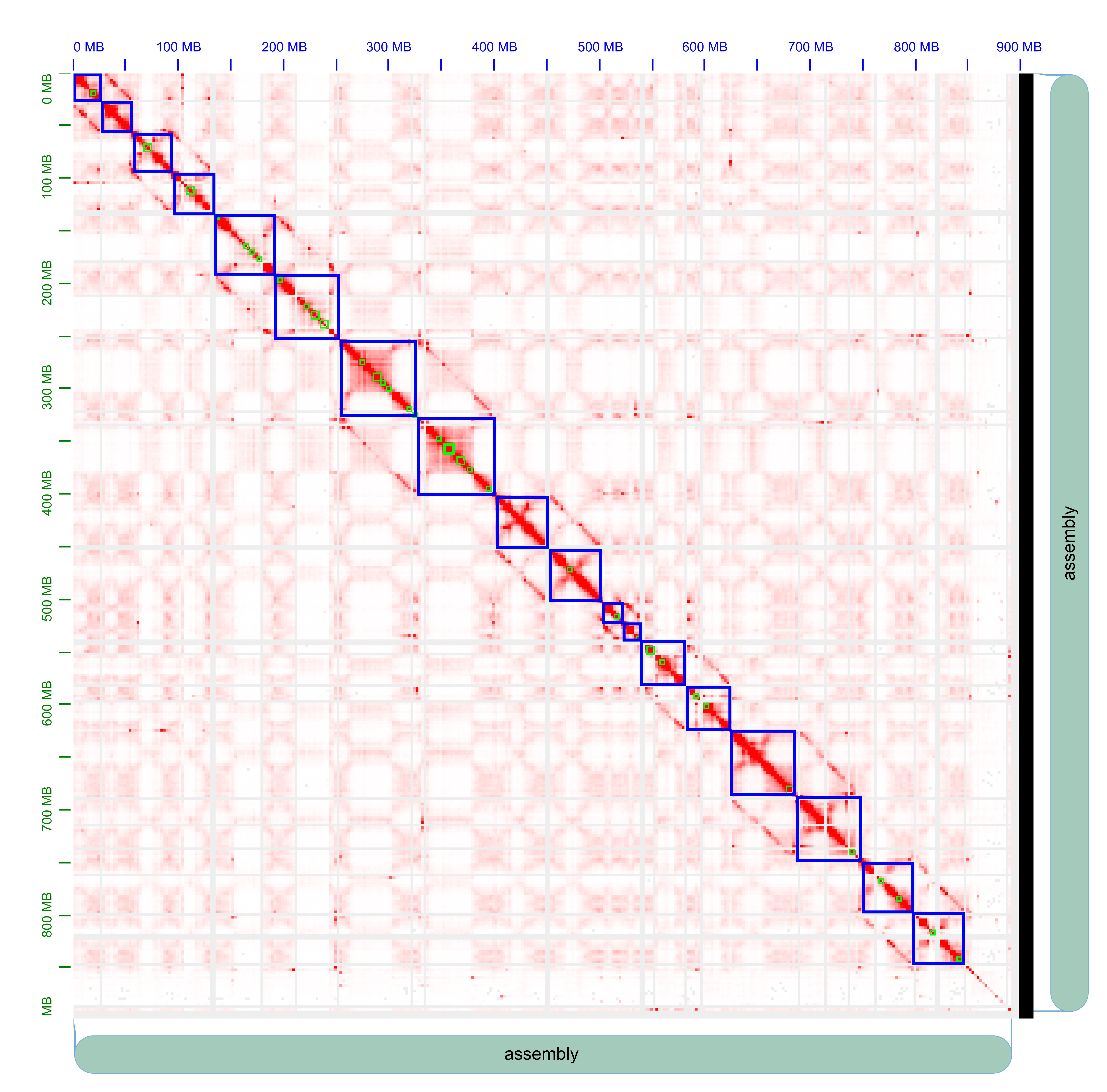

### Supplemental Figure 4

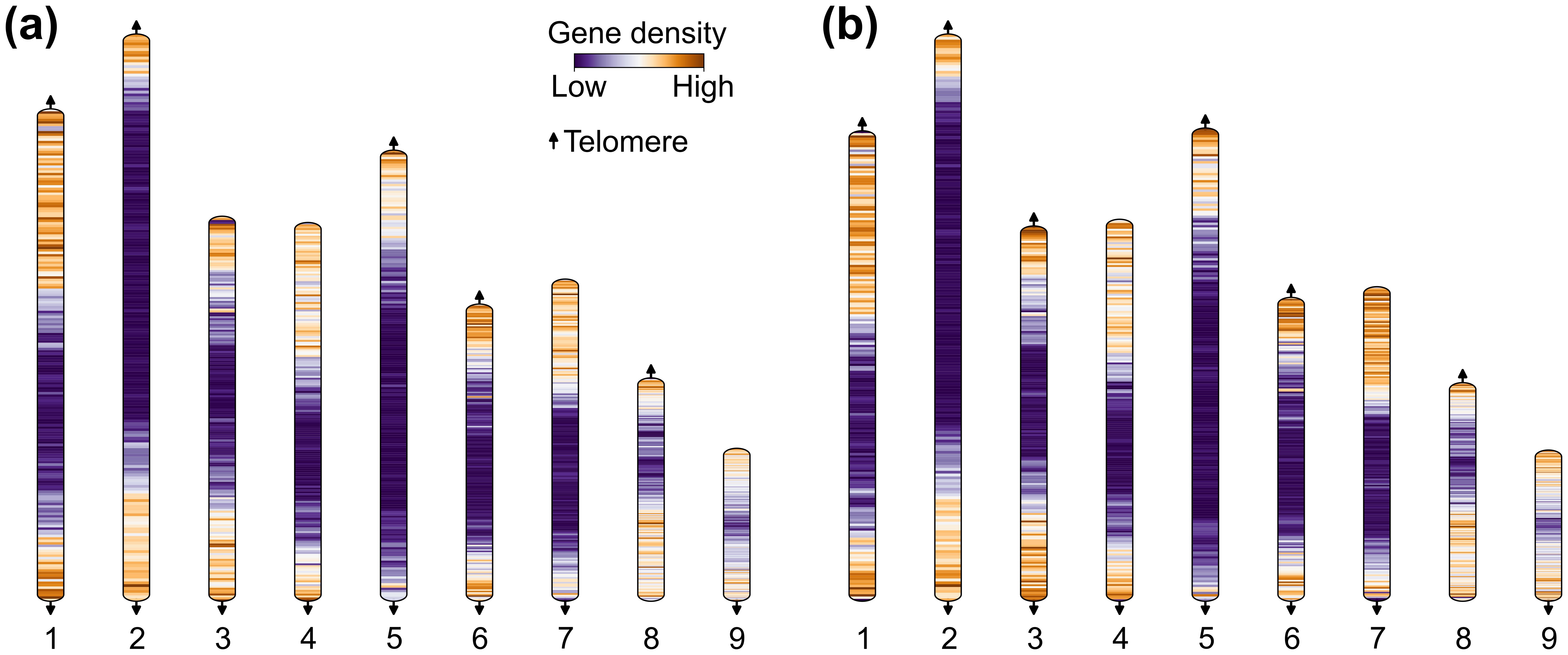
